# Tailoring the assembly of collagen fibers in alginate microspheres

**DOI:** 10.1101/2020.09.23.309823

**Authors:** Sarah Lehnert, Pawel Sikorski

**Affiliations:** Department of Physics, Norwegian University of Science and Technology (NTNU), Trondheim, Norway

**Keywords:** Collagen fibrillogenesis, Alginate, Microspheres, Tissue Engineering

## Abstract

The application of microspheres instead of bulk hydrogels in cell-laden biomaterials offers multiple advantages such as a high surface-to-volume-ratio and, consequently, a better nutrition and oxygen transfer to and from cells. The preparation of inert alginate microspheres is facile, quick, and well-established and the fabrication of alginate-collagen microspheres has been previously reported. However, no detailed characterization of the collagen fibrillogenesis in the alginate matrix is available. We use second-harmonic imaging microscopy reflection microscopy and turbidity assay to study assembly of collagen in alginate microspheres. We show that the assembly of collagen fibers in a gelled alginate matrix is a complex process that can be aided by addition of small polar molecules, such as glycine and by a careful selection of the gelling buffer used to prepare alginate hydrogels.

**Highlights:**

- *In situ* characterization of collagen fiber assembly in a gelled alginate matrix using collagen-specific second harmonic generated microscopy
- Collagen fibrillogenesis is positively influenced by the presence of small molecules in the solution prior microsphere preparation
- The ratio and amount of calcium, sodium and chloride ions used for the alginate gelling has also a crucial impact on the development of a collagen fiber network

## 1 Introduction

Tissue engineering (TE) is a promising research approach for the replacement or regeneration of damaged tissue *in vivo*. The focus lies on either implanting scaffolds, unseeded or seeded with cells, directly in or close to the damaged tissue to set off a regeneration cascade or to create living, functioning tissue *in vitro*. The created tissue can either be used for future *in vivo* transplantation or as a functional 3D-model for research purposes. Cell-based tissue engineering involves the culture of tissue specific cells or cell lines, and their seeding in or on three-dimensional scaffolds. The number of approaches to mimic natural tissue are as large as there are different tissues in the body. One, widely used strategy is to use hydrogels which mimic certain aspects of the natural microenvironment that cells experience *in vivo*. Hydrogels are described as three-dimensional, polymeric networks which are chemically or physically crosslinked. [1] They are capable of retaining a large amount of water and are therefore a good candidate to mimic the high-water content of the extracellular matrix (ECM) of cells *in vitro*. [2] The polymers used for such hydrophilic networks can be of natural or synthetic origin. A main consideration for choosing a hydrogel for TE applications is the ability of the chosen polymer to mimic the *in vivo* ECM of cells as closely as possible *in vitro* to induce *in vivo*-cell behavior such as proliferation and differentiation.

The *in vivo* ECM of cells is mainly comprised of collagen and elastin fibers, together with various polysaccharides and glycoproteins. The composition varies depending on the tissue. For instance, the ECM of bone cells consists to 60 % of inorganic, calcium-deficient hydroxyapatite and 40 % organic compounds. Furthermore, these organic compounds are to 90 % comprised of collagen type I and 10% of other proteins. [3] The protein collagen is comprised of three, situated parallel to each other, polypeptide strands. These strands consist of a repetitive X_aa_Y_aa_Gly-amino acid sequence. Owing to the glycine molecule at every third spot in each polypeptide strand, the single strands form ultimately a helical conformation coil with each other, intertwining to a right-handed triple helix. [4] Several methods can be used to evaluate the structure, orientation, and overall distribution of collagen fibers in biological samples. Besides FT-IRIS [5,6] and MRI [7,8], multiphoton-based microscopy (MPM) is widely used. [9–11] MPM is based on the nonlinear interactions between matter and photons. In conventional confocal fluorescence microscopy, one photon is absorbed by a fluorophore which consequently emits a single fluorescent photon with a lower energy. In contrast, in MPM two (or more) photons in the near infrared range (λ>700nm) are absorbed simultaneously in a single event. Consequently, emitted fluorescence has the energy that is twice the energy of the excitation light. [12,13] Besides the usage of MPM for fluorescence microscopy, a large application field is the imaging of unlabeled tissue samples such as collagen fibers by a process called second harmonic generation. [14] Collagen molecules, with their’ unique, non-centrosymmetric, molecular structure are capable of interacting with two incident photons in the focal spot simultaneously, without energy absorption. Hence, there is no energy loss and the process emits a scattered photon at half the wavelength of the two interacting photons. [15] Another method for the detection of assembled collagen fibers is confocal reflection microscopy. Here, the intrinsic optical property of collagen to back-scatter incoming light is used to detect the presence and structure of collagen fibrils. [16,17] However, it has been reported that only collagen fibers aligned with the imaging plane are detected. [18] Therefore, the combination of second harmonic multiphoton and confocal reflection microscopy is a unique tool to evaluate collagen specimen *in situ* without any previous staining, sectioning, or chemical processing. While collagen is also a ubiquitous component of the extracellular matrix and therefore a polymer that favors cell attachment and survival, a hydrogel prepared from unmineralized collagen alone often lacks the necessary mechanical strength that it exhibits in their natural, *in vivo* environment due to the lack of appropriate covalent crosslinking. [19,20] Additionally, the preparation of complex 3D collagen hydrogels can be difficult and attempts to prepare collagen microspheres include several processing steps. [21,22] In order to increase the mechanical strength but still benefit from the advantages of collagen, additional components such as alginate[23,24] or chitosan[25] can be blended in to be incorporated in hybrid hydrogels with collagen.

Alginate occurs naturally in brown seaweed and is an anionic polymer at natural pH. It is of particular interest for tissue engineering applications because it is non-toxic and biocompatible. The polymer is comprised of two block copolymers, namely (1,4)-linked β-D-mannuronate (M) and α-L-guluronate (G) residues. The amount, location and alternating pattern of those residues in the linear copolymers vary between different sources and determine its’ properties. [26] Alginate forms a crosslinked hydrogel in the presence of divalent cations such as Ca^2+^, Ba^2+^ or Sr^2+^ which interact mainly with the G-residues block in the polymer chains. [27] Furthermore, the mechanical strength of such alginate hydrogels can be tailored by varying alginate type (G-content and distribution of G-units), gelling conditions and alginate concentrations. Alginate itself is an inert material which lacks native cell adhesion ligands. Hence, the incorporation of collagen fibers is a logical step towards creating a biomaterial which mimics the backbone of the *in vivo* cell environment. For instance, da Cunha *et al.* demonstrated the fabrication of a three-dimensional alginate-collagen interpenetrating network as a substrate for fibroblast adhesion and a possible wound healing application. [28] Most hydrogel-based approaches aim to fabricate a three-dimensional model in the range of milli- to centimeters. With those dimensions, a major bottleneck is the sufficient transport of nutrients and oxygen by passive diffusion. In order to fully maintain a cellular function, cells need to no farther than 200 um away from the next capillary or surface. [29,30] This is a limitation that is hard to resolve when using bulk hydrogels.

Several groups aimed to prepare alginate-collagen spheres in the past, however, they did not fully explore the characteristics and limitations of the fabrication process in terms of morphology of collagen fiber that assembled and how one can specifically tailor it. Several studies have been published that describe preparation of alginate-collagen microspheres for applications in cell encapsulation. Ali *et al*. prepared 2% alginate-0.2% collagen microspheres for cell-encapsulation using the coaxial air-jet method in a solution containing 2% CaCl_2_. The authors described various outcomes for parameters for the fabrication and resulting geometrical values for the spheres as well as cell distribution. This study focused on the characterization of the retainment of a pro-angiogenic (FGF-2) molecule within the structure of the microspheres and how the incorporated cells reacted. [31] Mahou *et al*. also used the coaxial air-jet method to prepare 1% alginate-0.5% collagen cell-containing microspheres in 90mM CaCl_2_ solution. This study had the purpose of preparing spheres for the in vivo transplantation in mice to investigate the potential for promoting blood vessel development[32]. Workman *et al.*, use an electrostatic droplet generator (Bio-electrospraying) to generate 1,5 % alginate-1% collagen microspheres in 100mM CaCl_2._ The influence of preparation parameters such as nozzle size, precursor solution flow rate and applied voltage has been thoroughly investigated in this study. [33]

In this work, we identify factors that can support or suppress collagen fibrillogenesis in alginate gels. First, we show how collagen fiber assembly in an alginate matrix can be controlled and enhanced by the addition of amino acids to the alginate-collagen solution prior to microsphere preparation. Secondly, we suggest that the structure of the gelled alginate matrix as well as the amount of calcium- and sodium chloride has a significant effect on the type and quantity of assembled collagen fibers. We use collagen-specific second harmonic generated, as well as confocal reflection microscopy to describe the assembly of collagen within the alginate microsphere matrix *in situ*. We conducted additional SEM imaging of several samples to support the hypotheses we made after evaluating our alginate-collagen-constructs.

## 2 Experimental

### 2.1 Chemicals

All chemicals have been purchased from Sigma Aldrich (Norway) unless stated otherwise. De-ionized milli-Q water (MQ-water, 18.2 MΩcm) has been used throughout the experiments. Sodium alginate (Protanal LF200S, M_W_ = 2.74 × 10^5^ g mol^−1^, fraction of G-monomers: F_G_ = 0.68, F_GG_ = 0.57 and F_GM_ = 0.11) has been used for the microsphere preparation. Stock solutions of 3 wt-% alginate in MQ-water have been prepared prior to the experiments.

Table 1 summarizes the compositions for different gelling buffers used in this study. Stock solutions of every gelling buffer has been buffered with 10 mM Tris(hydroxymethyl)aminomethane (TRIS) at pH 7.3.

**Table 1.** Composition of gelling buffer for microsphere preparation.

| Gelling buffer | CaCl<sub>2</sub> | NaCl |
| --- | --- | --- |
| GB (10,50) | 10 mM | 50 mM |
| GB (25,25) | 25 mM | 25 mM |
| GB (50,25) | 50 mM | 25 mM |
| GB (50,50) | 50 mM | 50 mM |
| GB (50,75) | 50 mM | 75 mM |
| GB (75, 75) | 75 mM | 75 mM |

### 2.2 Microsphere preparation

The final sample volume of 2mL contained 0.6 wt-% alginate and 0.07 wt-% collagen. For the preparation of the collagen precursor solution, Collagen type I from rat tail has been purchased from ThermoFisher and the manufacturers protocol has been altered by replacing the suggested 0.1M PBS buffer with 0.1M MOPS, pH 7.4 in the precursor solution. Phosphate ions are reported to directly interact with collagen in their fibrillogenesis [34], as well the spontaneous precipitation of calcium phosphates upon the contact with a CaCl_2_-containing solution.[35] The precursor solution contained 0.1 µM Phenol red, a pH sensitive dye to monitor the pH of the solution. The collagen precursor solution has been vortexed and left to homogenize at 4 °C for 30 minutes.

Figure 1 shows the experimental workflow for the microsphere preparation. The final precursor solution contained 0.6 wt-% alginate and 0.07 wt-% collagen. For experiments including the addition of amino acids, samples have been prepared by mixing the alginate component with the respective amino acid before adding the collagen precursor. Microspheres have been prepared by dripping the alginate-collagen solution into a gelling bath (see Table1) at a rate of 15mL/h, a nozzle diameter of 0.35 mm and a voltage of 7 kV.

**Figure 1.**
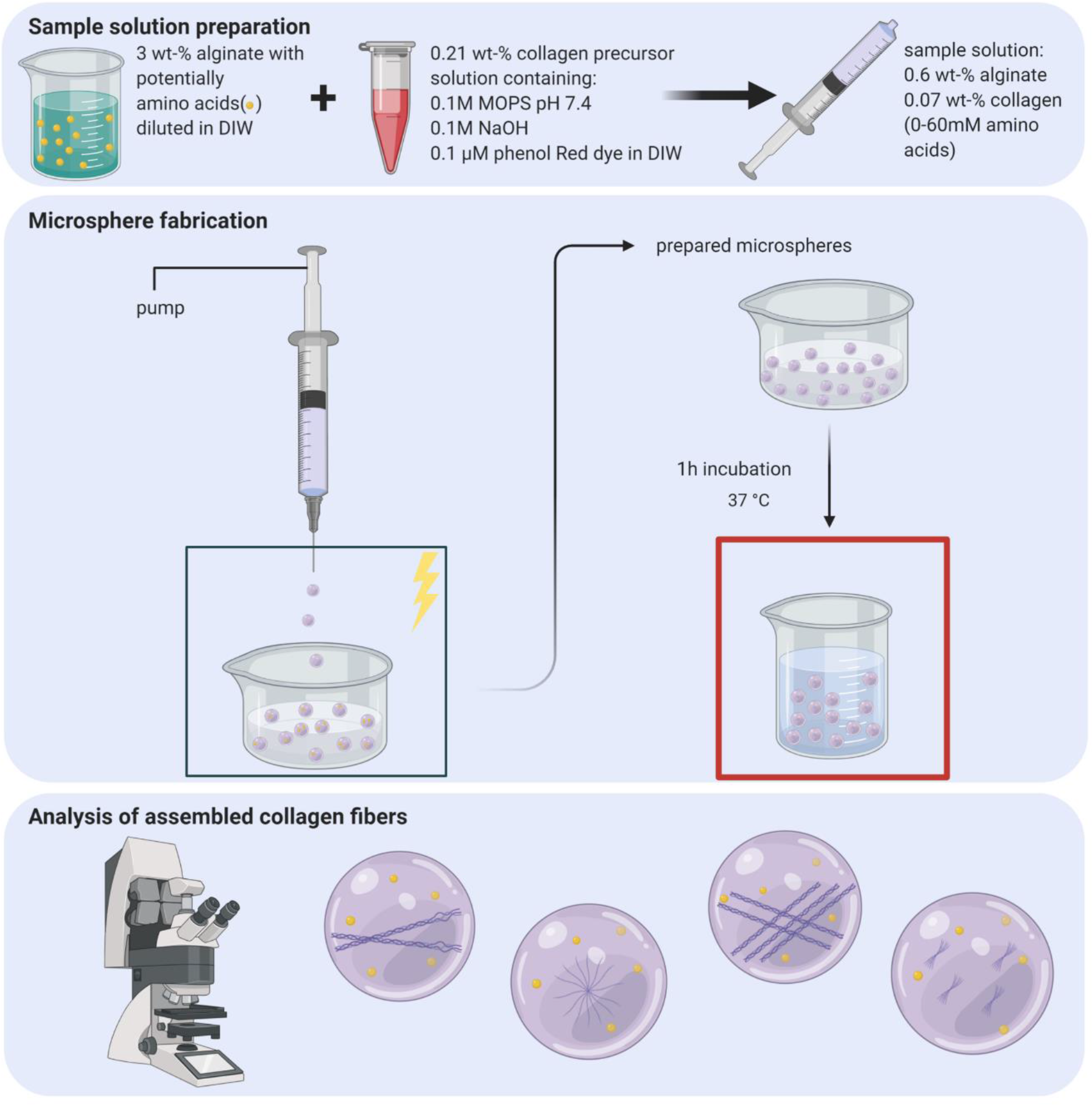
Schematic representation of the experimental workflow. The alginate solution and a collagen precursor solution were mixed in a syringe and the microspheres are prepared using an electrostatic droplet generator. The prepared microspheres were transferred to beakers containing fresh gelling buffer and incubated at 37 °C for 1h. The microspheres have been imaged with confocal microscopy.

The resulting microspheres have been collected from the gelling bath using a 150 µm polypropylene filter. In order to complete the gelling process, the sample has been placed in 5 mL of the respective gelling buffer and kept for 1h at 37 °C.

### 2.3 Preparation of bulk hydrogels

For the initial study regarding the likelihood of collagen fibers assembling in an 0.6 wt-% alginate matrix, bulk hydrogels have been prepared. A collagen precursor solution has been prepared as described previously and added to an alginate solution to a final concentration of 0.6 wt-% alginate and 0.07 wt-% collagen. 60 µL of the alginate-collagen solution has been cast in a silicon mold with a diameter of 8.5 mm and a height of 1.7 mm (Grace Bio Labs, press-to-seal silicon isolators) and placed in a petri dish. The mold and the gel have consequently been covered with the gelling buffer GB (50,50) (see Table 1) and incubated for 1h at 37 °C. The CLEX hydrogels have been prepared according Bassett *et al.* with a final alginate concentration of 0.4 wt-% and a collagen-concentration of 0.07 wt-%. [36] The concentration of glycine and ethylenediaminediacetic acid (EDDA) was in both systems 60 mM.

### 2.4 Alginate gelling kinetics and collagen fibrillogenesis kinetics

For the alginate gelling kinetic analysis, a previously established protocol by Bjørnøy *et al*. has been applied. [37] Briefly, a flow cell has been prepared by placing a 1.5 µL droplet of 0.6 wt-% alginate solution between a microscope slide and a thin cover slide. A single layer of a 140 µm thin double-sided tape has been used as a spacer. Applying gentle pressure to the setup compressed the alginate droplet into a disc shape. Gelling has been initiated by applying 50 µL of gelling buffer (Table 1) into the spacing between the two slides. The gelling has been recorded by using an Andor Zyla SCMOS camera attached to a Nikon Eclipse TS100 microscope. The progression of the gelling front through the alginate solution was monitored by using a 4x/0.13NA Nikon Plan Fluor phase contrast objective. For analyzing the resulting gelling videos, a macro for ImageJ has been developed which follows the progression of the gelling front with a line plot. After subtracting the background, the resulting data set has then been plotted using python and the slope of the linear region of the gelling step has been calculated.

A possible glycine-alginate interaction has been investigated twofold. For one, the gelling kinetics of 0.6 wt-% alginate containing 6mM glycine have been measured, similar to the method described above. Additionally, viscosity measurements have been conducted to detect possible a glycine-alginate interaction at the molecular level. Briefly, the viscosity of a 0.8 mg/mL alginate solution containing an increasing amount of either glycine, serine or arginine in 0.1M MOPS, pH 7.4 has been measured six-fold at room temperature with a Mikro-Ubbelohde-Viscosimeter and the relative viscosity has been calculated.

For determining the individual impact of gelling buffers, presence of amino acids and ungelled alginate on the assembly of collagen fibers, the change in absorbance of 0.12 wt% collagen at 350 nm and at 37 °C has been measured using a the SpectraMax® i3 platform spectrophotometer every 35 seconds for one hour. The collagen precursor solutions have been prepared like the precursor solutions for the microsphere preparation but at a slightly higher concentration of 0.25 wt%. In a second step, the precursor was mixed with MQ-water with or without amino acids. Finally, this sample solution has been distributed between several tubes containing the different gelling buffers. After thorough vortexing, each collagen-solution-sample has been added to three wells in a 96-wellplate and consequently measured.

### 2.5 Evaluation of collagen fiber assembly using reflection confocal Microscopy (RCM) and Second-harmonic Imaging Microscopy (SHIM)

Alginate-collagen microsphere samples were transferred to 12-well glass bottom plates (#1.5 High Performance cover glass, Cellvis) and mounted in a confocal microscope TSC SP8 MP(Leica). The microscope is equipped with standard photomultiplier tubes (PMT’s) and, more sensitive, hybrid detectors (HyD’s). As light source, a white light laser and a tunable, near-infrared laser Chameleon Vision-S (Coherent) has been used. The respective samples have been imaged with a 25x water objective, NA=0.95. The reflection signal of collagen has been measured in the range of 475-495 nm with an internal PMT. The sample was illuminated with incident light at 780 nm and the reflected SHG signal was detected using a HyD detector without pinhole in the range of 370 – 410 nm. The laser power was minimized to prevent sample damage. All images have been taken with a frame size of 1024×1024 and an imaging speed of 100 Hz. For signal enhancement, the SHIM signal has been measured with a line accumulation of 16 and the reflection signal with a line average of 5.

The microscope has a known offset of 5 um between the internal detector for the reflection and the external detector for the SHIM signal. This prohibits the overlay of the reflection and SHIM signals. Therefore, the images have been divided in a SHIM-signal and a reflection signal part. For the quantitative analysis of the assembled collagen fibers, SHIM images have been analyzed with ImageJ. After setting an intensity threshold to exclude background noise, particles greater than 1.5 µm have been detected using the analyze particle algorithm, resulting in an overview of the total count of individual particles and their respective area in µm^2^. In total, three representative SHIM images for each sample have been analyzed.

### 2.6 Sample preparation for scanning electron microscopy (SEM)

Microspheres prepared as previously described in either GB 10,50 or GB 50,75 without additional amino acids have been dried using a critical point dryer (CPD; Quorum K850). The respective collagen gels for a further analysis of the collagen fibrils have been prepared on a soft silicon substrate, similar to the described preparation of bulk hydrogels, to prevent the collagen assembly on a hard surface, left in the oven for one day and then consequently dried with the CPD. The dried samples were collected and mounted on SEM stubs using liquid silver paint (Pelco®) to increase conductivity. Prior imaging with a SEM APREO system, the samples have been coated with 15 nm platin/palladium (Pt/Pd 80/20) and imaging has been conducted at 1 kV, 6.3 pA and an additional threshold set to 50 V to prevent surface charging due to the high porosity of the samples as well as damaging of the surface.

For the analysis of the inner alginate-collagen structure of the microspheres prepared with GB (10,50) and GB (50,75), the respective samples have been embedded in 2 wt-% agarose and cut in slices using a Vibratome VT1000S from Leica prior the CPD-drying.

## 3 Results

### 3.1 Studies from CLEX-alginate-collagen-hydrogel preparation provide clues on how to improve fibrillogenesis of collagen in alginate microspheres

The aim of this study is to prepare an interpenetrating alginate-collagen network (IPN). In such IPN, the collagen has several functions. It could support cell attachment and serves as scaffold for enzymatically induced mineralization using alkaline phosphatase (ALP). Alginate-collagen IPN were prepared using rat tail collagen type I, that is often used to prepare hydrogel scaffolds for studies of cellular processes in 3D. [28,38,39]

Before the preparation of the IPNs, the pH of the collagen precursor solution was increased according to the protocol used to make 3D-collagen hydrogels (see Experimental). The change in pH induces collagen fiber assembly, a process that takes about 40 minutes at 37 °C. [40] Neutralized collagen precursor solution was added to an alginate solution to achieve a collagen concentration of 0.07 wt-% and an alginate concentration of 0.6 wt-%. Alginate-collagen microspheres were made with an electrostatic droplet generator (see Figure 1). Prepared microspheres have been characterized by laser scanning confocal microscopy using second-harmonic imaging microscopy (SHIM) and reflection confocal microscopy (RCM) modalities (see Experimental). In SHIM, the signal is generated by a nonlinear optical process known as second-harmonic generation (SHG). In our composites, a SHIM signal can only be generated by correctly assembled collagen fibers. [9,11] Reflection confocal microscopy is based on the ability of collagen fibers to reflect and scatter incoming laser light. The signal is generated at the interface between hydrogel regions with differences in the refractive index. [41] All microsphere samples have been imaged at their respective equatorial plane to depict a cross-section throughout the whole microsphere.

Figure 2A shows a brightfield image of the prepared microspheres made in a gelling bath containing 50mM CaCl_2_, 50mM NaCl and 10mM TRIS with a pH of 7.3. The respective SHIM and reflection signals for the microspheres is shown in Figure 2B. Both, RCM and SHIM signals, are weak, indicating poorly developed collagen fiber network. In contrast, the collagen assembly in a larger bulk gel, containing the same concentration of alginate and collagen and prepared using the diffusion method (see Experimental), resulted in a much higher signals from collagen fiber bundles (see Figure 2C). Based on this observation, we can hypothesize that it is possible to generate alginate-collagen IPNs in which a fibrillar collagen network is formed. Additionally, we prepared bulk hydrogels using the CLEX method. [36] In these gels, the gelling is driven by a competitive ligand exchange of crosslinking ions in the presence of chelators such as ethylenediaminediacetic acid (EDDA) or glycine. Figure 2D and 2E show the collagen fiber networks for collagen-alginate IPNs prepared in the presence of EDDA and glycine using the CLEX approach. The collagen fibers in the hydrogel containing EDDA (Figure 2D) assemble to a uniform network of thin fibers with a diameter of roughly 2 µm. In contrast, in the presence of glycine, longer and thicker collagen fibers (diameter of 4 µm) in a finer network of thinner fibrils (diameter of 0.5 µm) were observed (Figure 2E). This results in strong SHIM (yellow) and reflection (cyan) signals in CLSM. We therefore hypothesized that certain additives might improve collagen assembly in alginate-collagen IPNs.

**Figure 2.**
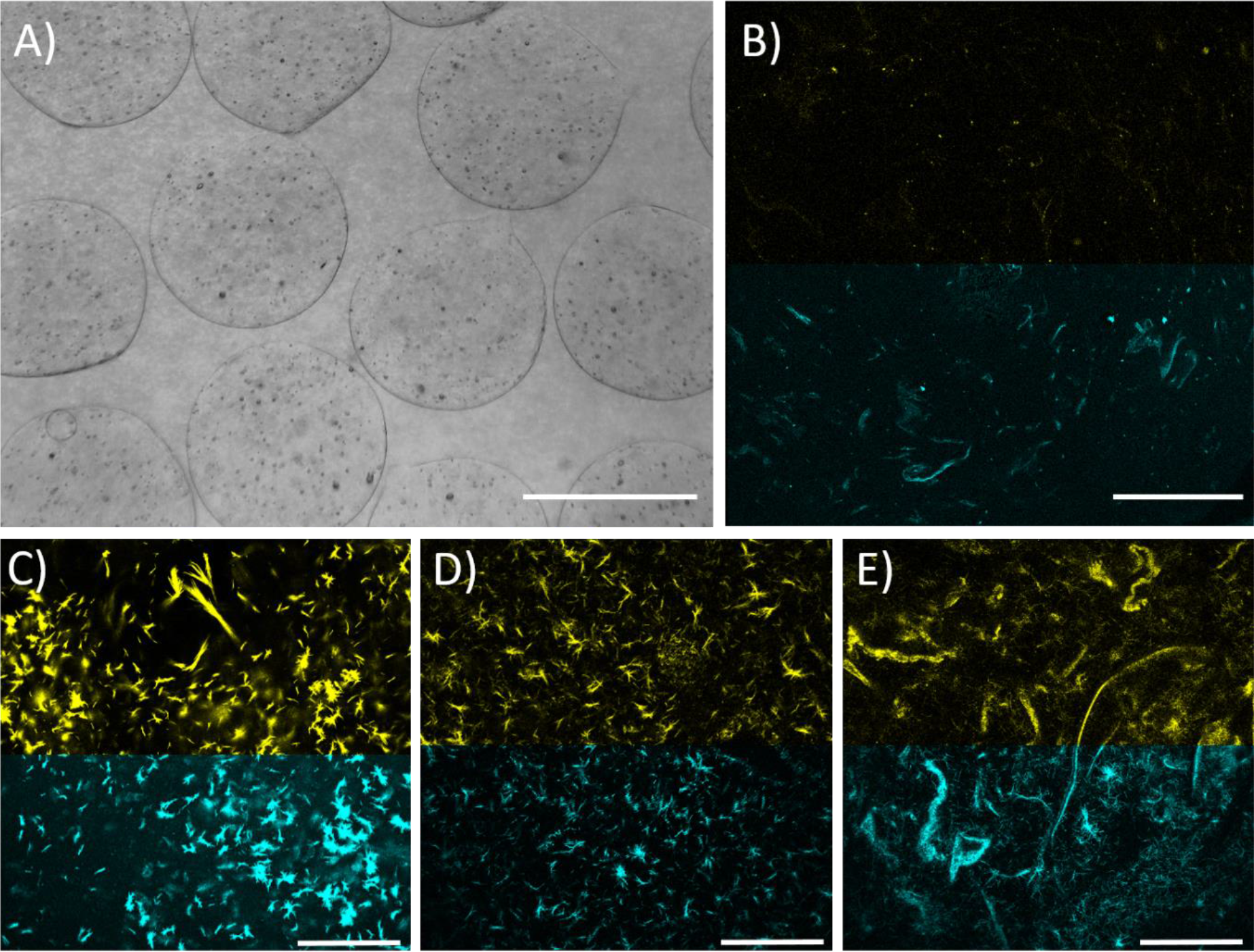
Assembly of collagen fibers in alginate microspheres. Brightfield image of microspheres prepared in gelling buffer containing 50mM CaCl2 and 50 mM NaCl is shown in A (Scale Bar: 600 um). Corresponding SHIM and RCM signals for these microspheres are shown in B. Little assembled collagen can be detected for this composite. C) Bulk gel prepared at the same conditions, shows well developed collagen network that is detected both with RM (cyan) and SHIM (yellow). Hydrogels prepared using the CLEX method using EDDA or Glycine as chelating agent are shown in D and E respectively. Collagen network is detected in these gels with RCM and SHIM. Scale bar: 150 µm.

### 3.2 The addition of glycine, serine, proline and arginine to the alginate-collagen solution influences the overall collagen fibrillogenesis process

As shown in Figure 2D and 2E, the assembly of collagen in an alginate matrix can be enhanced and altered by the presence of EDDA and glycine. To confirm this observation, microspheres with an increasing concentration of glycine (0.01 mM to 60 mM) in the alginate-collagen solution were prepared. This concentration series visualizes the effect that glycine has on the collagen fiber assembly (Figure 3A). With an increasing amount of glycine in the alginate-collagen solution, the amount of collagen fibers detected with both, reflection and SHIM was increased. A corresponding quantitative analysis is presented in Figure 3B. The fabrication of alginate-collagen microspheres without additional glycine in the solution (0mM; Figure 3A-i), let to the assembly of only a low number of collagen fiber bundles with a small area size compared to the samples containing additional glycine. Both, the total collagen area throughout the imaged microsphere cross-section as well as the number of individual collagen fibers increases with an increasing amount of glycine in the alginate-collagen solution prior microsphere preparation.

**Figure 3.**
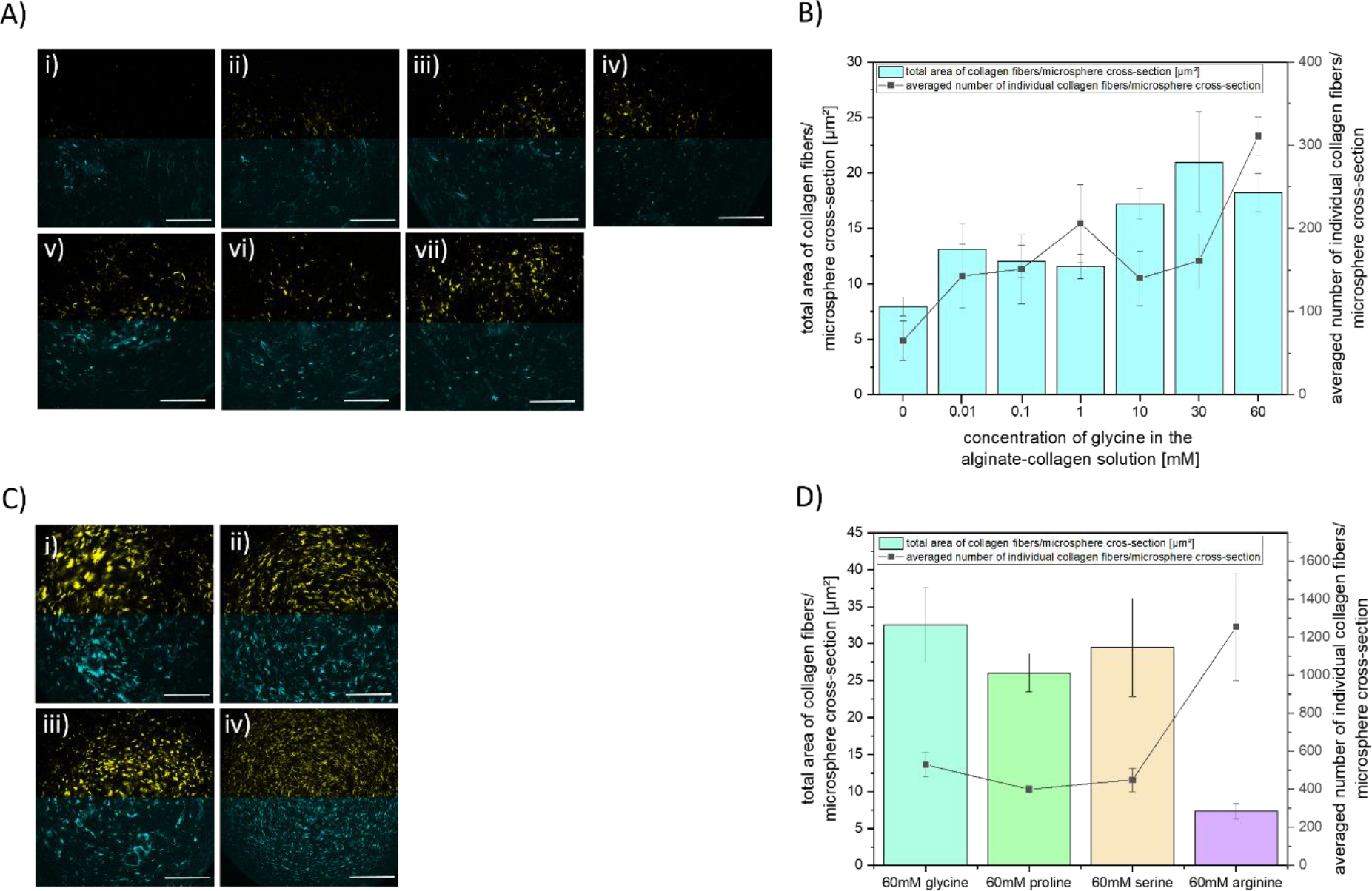
Effect of the addition of amino acids on the assembly of collagen fibers within an alginate microsphere-matrix. All samples have been gelled in a gelling bath containing GB (50,50). A) Confocal reflection and SHIM signals for alginate-collagen microspheres prepared without glycine (i) and with an increasing concentration of glycine of 0.01mM (ii), 0.1mM (iii), 1mM (iv), 10mM (v), 30mM (vi) and 60mM (vii). Yellow: SHIM-signal. Cyan: reflection signal. B) Collagen fiber analysis for the samples containing an increasing amount of glycine. Bars represent the total area of detected collagen fibers per respective microsphere cross-section, and the line shows the number of separately detected collagen fibers in the imaged microsphere cross-section. SHIM images from 3 microspheres have been analyzed for each condition. Effect of 60mM glycine (i), proline (ii), serine (iii) and 60mM arginine (iv) on the collagen fiber assembly (C) and corresponding area and number of detected collagen fibers (D). Scale bar: 150 µm.

To assess if the observed effect is due to specific interactions between glycine and collagen or glycine and alginate, microspheres were prepared with 60 mM proline, serine and arginine. In addition, the phenol red included in the collagen precursor solution (see Experimental section) was used to monitor the pH of the alginate-collagen solution. The addition of glycine, serine and proline respectively did not alter the overall pH of the solution and confocal microscopy images for those microspheres are shown in Figure 3C. Similar assembly of collagen fiber bundles was observed in these samples with glycine giving the strongest overall signal (Figure 3C-i). The situation was different for the arginine-containing microspheres. The pH of the alginate-collagen solution increased once the arginine was added and this was observed by a color change of the solution from orange to pink. The confocal analysis showed that both, reflection and SHIM signal, displayed a much finer fiber structure of assembled collagen (Figure 3C-iv). A particle analysis of the samples prepared with 60mM of different amino acids is shown in Figure 3D. Depicted is the averaged area of detected collagen fibers in the imaged cross-section (in µm^2^) as well as the average number of separately detected collagen fibers in the same imaging plane. The overall area that is made up of assembled collagen fiber bundles in the microsphere cross-section as well as the number of detected collagen fibers is similar for glycine, proline and serine, whilst the sample prepared with 60mM arginine contained roughly three-times the amount of separate, but small, collagen fiber bundles.

### 3.3 The ratio of CaCl_2_ and NaCl in the alginate gelling buffer during the preparation and incubation step is important for a successful assembly of collagen fibers within the alginate matrix

After determining a favorable role of additives on the assembly of collagen fibers in a gelled alginate matrix, the question remained to what extent not only the gelling process but also the individual components of the gelling buffer influence the collagen fiber assembly. To answer this question, six different gelling buffers were prepared. Three of them contained equal molar concentrations of CaCl_2_ and NaCl, while the other three contained different ratios of CaCl_2_ to NaCl. Up to this point, the microspheres, and hydrogels for Figures 2 and 3 have been prepared using a gelling buffer containing 50mM CaCl_2_, 50mM NaCl and 10mM TRIS with a pH of 7.3. The nomenclature for the different gelling buffers is as followed, the concentrations of CaCl_2_ and NaCl are given in mM and put in brackets. Every gelling buffer additionally contained 10mM TRIS and the pH was set to 7.3. Microspheres of 0.6 wt-% alginate and 0.07 wt-% collagen have been prepared in MQ-water. The alginate-collagen solutions were prepared with or without 6mM glycine to further investigate its’ effect on the collagen assembly.

Confocal images for microspheres prepared in different gelling buffers are shown in Figures 4A and 4B. The only gelling buffer which resulted in a successful formation of a collagen fiber network in the absence of glycine was GB (50,75) (Figure 4B-v), and no assembled collagen was detected for the other gelling buffers. Microspheres prepared with 6mM glycine in the alginate-collagen solution and in gelling buffers containing 50mM CaCl_2_ (GB(50,50), Figure 4A-iv; GB(50,25), Figure 4B-iv; GB(50,75), Figure 4B-vi) shows an analogical assembly of collagen fibers in thick, short bundles. As for GB (50,75), the signal intensity for the collagen fiber bundles becomes stronger and suggested an even more interconnected network compared to the microspheres prepared without glycine for the same buffer. In the presence of glycine, using gelling buffer GB (25,25) for the microsphere preparation also resulted in assembled collagen fibers. However, the SHIM signal suggests the presence of very small collagen-particles which are not connected to each other. The reflection signal does show a few larger, fiber-like structures but those do not give a corresponding SHIM signal. Microspheres prepared with 6mM glycine in GB (75,75), Figure 4A-vi do not show the assembly of collagen fibers. A similar observation has been made for the gelling buffer GB (10,50), Figure 4B-ii.

**Figure 4.**
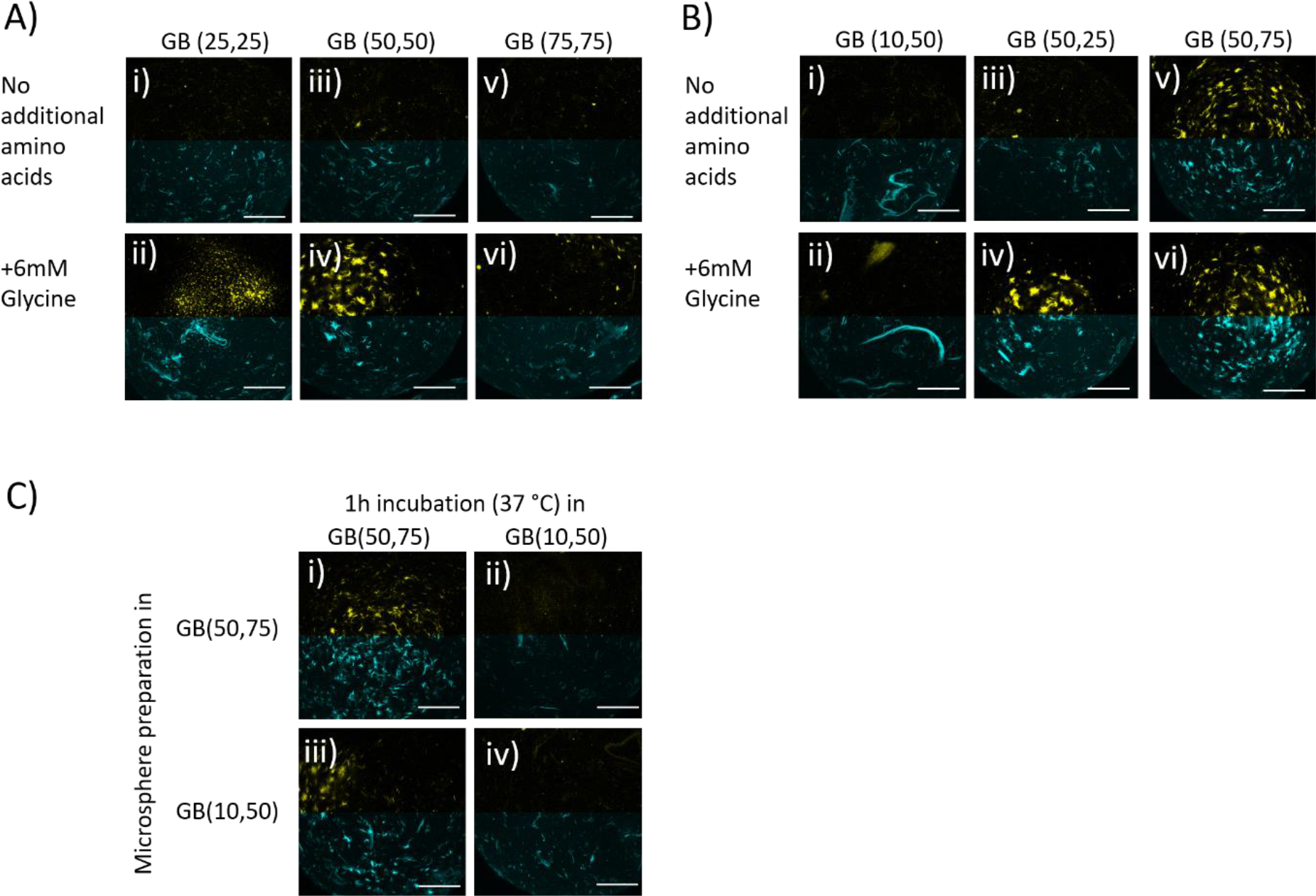
Influence of the gelling buffer used for the microsphere preparation on collagen assembly. A) Confocal analysis of assembled collagen fibers in alginate matrix using gelling buffers with an equimolar ratio of CaCl_2_ and NaCl. B) Confocal microscopy analysis of assembled collagen fibers in alginate matrix using gelling buffers with a different ratio of CaCl_2_ to NaCl. C) Effect of the gelling buffer for the microsphere preparation compared to the effect of the gelling buffer used for the 1h incubation at 37 °C. Samples shown in the top row have been prepared using GB (50,75) and samples in the bottom row have been prepared using GB (10,50). Collagen assembled only in sample shown in (i) and (ii) that after preparation have been incubated in GB (50,75).

It is interesting to note on what time scale collagen assembles and how this process seems to be affected by the used gelling buffer. By deliberately switching between preparation and incubation buffer, we were able to demonstrate that the choice of incubation buffer has the largest effect on the successful development of an IPN in the microspheres (Figure 4C). Additional surface observations conducted with SEM showed that the samples prepared and incubated in the same gelling buffer GB (10,50) and GB (50,75) respectively displayed significantly different characteristics in terms of porosity and homogeneity (Supplementary, Figure S1A and S1B) at both, the outer surface as well as in the middle of the microsphere.

### 3.4 Multiple factors such as CaCl_2_: NaCl ratio and alginate concentration have a direct influence on the kinetic of collagen fiber assembly

To fully understand the specific requirements under which collagen fibers assemble in an alginate matrix, two aspect of the IPN preparation have been investigated further. First, the speed of the alginate gelling has been determined for the gelling buffers used above. A small droplet of 0.6wt-% alginate has been placed between two glass slides and the advancing gelling front has been recorded with a microscope camera. [37] Measured gel front velocities are shown in Figure S5A (Supplementary). The velocity increases for higher Ca^2+^-concentrations and is between 1 and 3 µm/s. Furthermore, the addition of 6mM glycine to the alginate solution did not change the gelling speed significantly. Additionally, the possible influence of glycine on the alginate solution has been considered and investigated thoroughly by measuring viscosity of the alginate solutions containing a range of different glycine concentrations, the viscosity did not change (data not shown). We therefore concluded that the presence of amino acids in the solution has an impact on the collagen fiber assembly and does not influence the alginate gelling process.

Next, the influence of the gelling buffer composition on the collagen fiber assembly in the absence of alginate has been investigated using a turbidity assay (see Experimental). Turbidity measurements are a well-established method to follow the assembly kinetics of collagen fibers [42] by measuring changes in optical density during collagen fiber assembly which follows a sigmoidal curve. The impact of additional chemicals on the collagen fiber assembly can be directly followed by evaluating the changes of this sigmoidal shape in terms of lag-, growth- and plateau phases. The collagen precursor solution has been prepared in a similar way as the solution used to make alginate-collagen microspheres, but with an increased collagen concentration (0.12wt-%). The impact of separate NaCl- and CaCl_2_-solutions on the collagen assembly has been reported before [43] and by confirming these measurements (Supplementary, Figure S2A), we positively validated our analysis method. Briefly, collagen assembled in NaCl-solutions prior the measurement had been started and with an increasing CaCl_2_-concentration, the lag-phase of collagen assembly is significantly prolongated (Supplementary, Figure S2B and S2C). The combined effect of both, CaCl_2_ and NaCl, in the gelling buffers used to prepare the alginate-collagen microspheres in this study is shown in Figure 5A. The collagen in the control in 10mM TRIS, pH 7.3 assembled to a small amount before the measurement has been started, also in GB (10,50) and GB (25,25) the assembly started just after mixing. Samples containing either 50mM or 75mM CaCl_2_ displayed a significant lag phase prior the fiber assembly. The addition of NaCl shortens the lag phase (compare GB (50,75) and GB (50,25)), however, the final OD_350nm_ of those samples are similar, indicating that a similar collagen network has assembled. The preparation and analysis of corresponding collagen hydrogels further showed the assembly of collagen fibers with different thicknesses and, length and orientation (Supplementary, Figure S3). These findings suggest that the gelling buffer not only controls the velocity of the alginate gelling but also actively promotes or attenuates the collagen fiber assembly.

**Figure 5.**
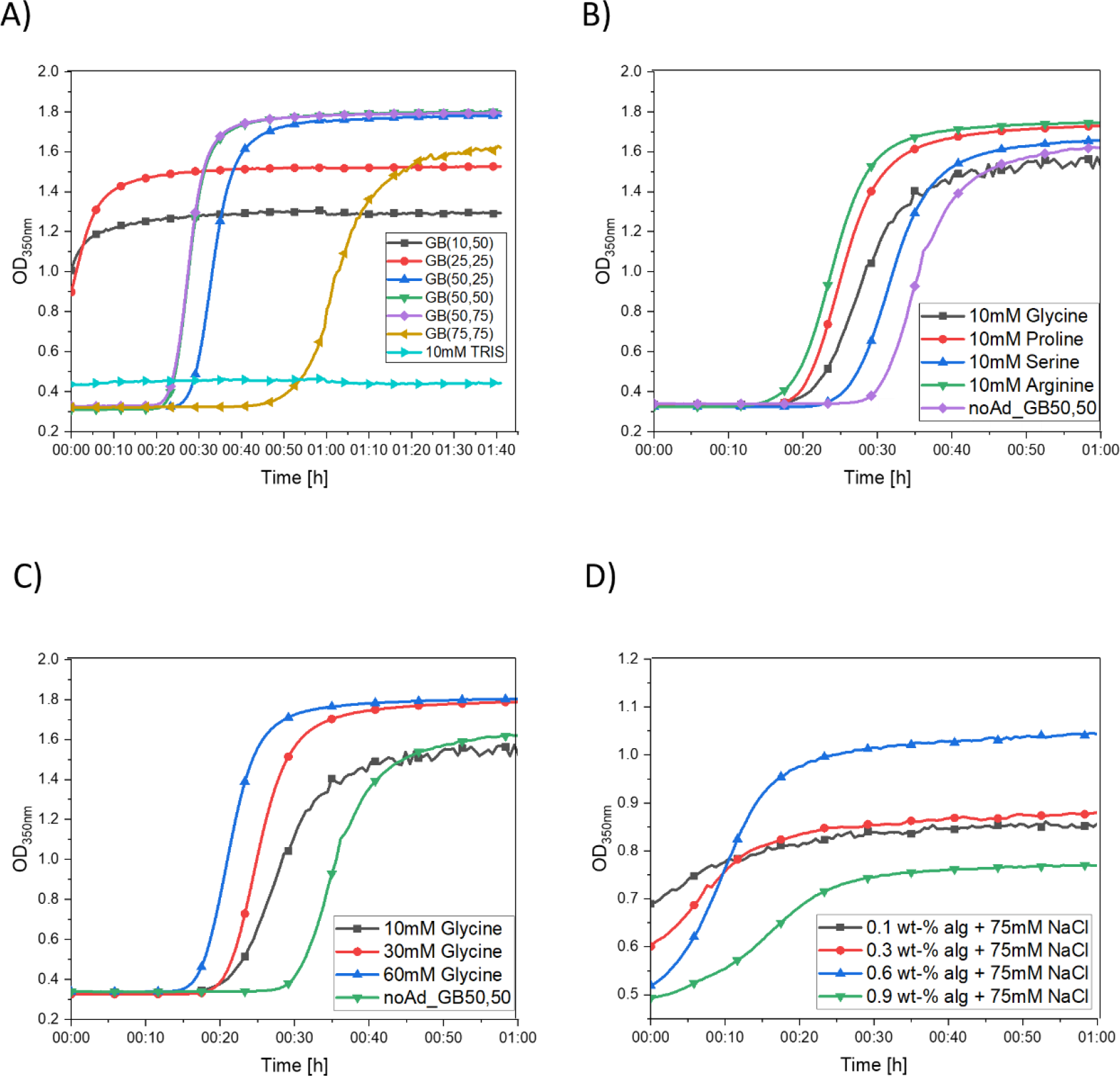
Collagen fibrillogenesis investigated at 37 °C using using turbidity assay. A) Influence of the gelling buffers used in this study on the collagen fiber assembly. B) Effect of amino acids with a concentration of 10 mM on the collagen assembly in GB(50,50). C) Effect of glycine on the collagen fiber assembly in GB(50,50) investigated as a function of concentration. D) Effect of ungelled alginate on assembly of 0.12 wt-% collagen.

The effect amino acids have on the collagen fiber assembly is shown in Figure 5B. All amino acids shorten the lag phase of the collagen assembly. Corresponding SEM analysis revealed a similar fiber architecture for collagen gels containing 10mM glycine, proline and serine, while the gel containing 10 mM arginine showed an increased number of collagen fibers which were also thinner (Figure S4). This in in good consensus with the observations made in Figure 3C and 3D, showing a similar assembly of collagen fibers in the alginate matrix for additional glycine, proline and serine but a vastly different collagen fiber morphology for arginine. An increase in concentration of the amino acids also has an effect on shortening the initial lag phase (Figure 5C), indicating that the concentration series made with glycine in Figure 3A and 3B does, indeed, yield in a higher number of assembled collagen fiber bundles.

Lastly, also the effect of alginate on the kinetics of collagen fiber assembly was investigated. The effect of molecular crowding of ungelled alginate molecules on the assembly of collagen fibers can be seen in Figure 5D. A concentration too low (0.1 wt-% and 0.3 wt-%) or too high (0.9 wt-%) suppresses the assembly of collagen fibers while a concentration of 0.6 wt-% alginate slightly delays but supports the collagen fibrillogenesis, indicating that the presence of alginate as an anionic polysaccharide approaches an optimum-concentration at which it supports the collagen fiber assembly.

## 4 Discussion

The presented results reveal that the fibrillogenesis of collagen in a gelling alginate matrix is a complex mechanism. The initial experiments in Figure 2 showed that the preparation protocol for bulk hydrogels cannot be easily transferred to the preparation of microspheres consisting of the same components. Collagen hydrogels are generally incubated for 40 minutes or longer to ensure fibrillogenesis. Based on the study of Bassett *et al.*, the alginate in the CLEX hydrogels gelled between 7-30 seconds. [36] This means that the detected collagen fibers in both hydrogels assembled despite a quickly gelled alginate matrix. Furthermore, the morphology of the collagen fibers varies significantly for the two CLEX hydrogels which only differ in the presence of either EDDA (Figure 2D) or glycine (Figure 2E). Physical parameters of the microsphere preparation procedure, such as applied voltage, nozzle diameter or pump speed, have been kept constant in this work but could also offer a subject for further optimization.

### 4.1 Influence of several amino acids on the collagen fibrillogenesis in a gelling alginate matrix

During the preparation of alginate-collagen microspheres containing an increasing amount of glycine, we observed an increase in number and size for detectable collagen fibers (Figure 3A and 3B). The determination of the gelling speed of a glycine-containing alginate solution as well as a viscosity study on the effect of glycine on the alginate solution revealed no changes compared to the pure alginate solutions used as controls. We consequently concluded that an interaction between the glycine molecules and the collagen molecules during its’ fibrillogenesis is likely. Investigating the same setup used for the preparation for Figure 3A but without the alginate-gelling component, we were able to verify that the addition of glycine in an increasing concentration to the collagen solution significantly shortens the initial lag-phase of the collagen fiber assembly (Figure 5C). Additionally, the higher the added glycine concentration, the higher was the overall OD which is generally understood as a sign for a higher amount of assembled collagen fibers. The effect of the addition of a separate solution of glycine to the *in vitro* fibrillogenesis of collagen has not been reported yet. In fact, only one group investigated the presence of molecules like lysine and glutamic acid on the *in vitro* collagen assembly. In contrast to our findings, they determined a prolongated lag-phase upon the addition of an increasing amount of lysine. [44] However, our system varies greatly from Liu *et al*.’s such as they raised the pH of the collagen solution by using 10mM sodium phosphate solution, pH 7.2 and the final collagen solution contained a very high amount of NaCl (110mM). Phosphate ions are known to interfere with the collagen assembly by building salt bridges. [34] Furthermore, the calcium present in the gelling bath to induce the gelling of the alginate microspheres would most likely interact with the phosphate ions and form calcium phosphate precipitates [45], making the characterization and analysis of the collagen fiber assembly more unpredictable and less controllable. Therefore, in our system, we avoided using known initiators or interfering agents of collagen fibrillogenesis and alginate gelling.

We discovered that the presence of different amino acid shortened the lag phase of the collagen assembly to different degrees (Figure 5B). While the exact mechanism is not entirely understood yet, it is interesting to note that the ability to shorten the lag phase is not dependent on the size of the molecule. Glycine is smaller than serine (75 Da compared to 105 Da) but initiates the growth phase faster. Arginine, which exhibits similar characteristics compared to the lysine used by Liu *et al.* (in terms of a positively charged side chain), displayed the shortest lag phase with cutting the time until collagen started to assemble nearly in half. The origin of the interaction of the separate amino acid solutions and the collagen molecules prior and during fibrillogenesis is still not clear. Derived from measuring the zeta potential, it has been reported that charge-interactions between the separately added and positively charged lysine and the collagen molecules take place. [44] Objectively comparing the different solutions we used at a neutral pH of 7.4, the molecules display different charges. Based on the isoelectric point (pI), serine (5.68), glycine (6.06) and proline (6.3) should have a negative charge, while arginine (10.76) will most likely be positively charged. Interestingly enough, the order of amino acids which shorten the lag phase from most to least follows the same scheme, namely the positively charged arginine has the greatest effect while serine, with the lowest pI and negatively charged, has the least effect on initiating the collagen assembly quicker (Figure 5B).

### 4.2 Influences of the gelling system on the collagen fibrillogenesis in a gelling alginate matrix

Inhomogeneities in the gelled alginate matrix are promoted by a lower CaCl_2_ concentration since a low concentration of crosslinking Ca-ions gives the alginate more time to diffuse to the sphere-surface and hence increases the concentration of alginate close to the bead surface. [27] Skjåk-Bræk reported that this gradient can be attenuated by adding a higher concentration of sodium chloride. [46] Applying this knowledge to the gelling buffers used in this study, a more homogenous alginate network should have developed for 50mM CaCl_2_ and a higher NaCl concentration of 75mM compared to GB (10,50). It is possible that the morphology of the gelled alginate matrix, too, affects the formation of collagen fibrils. The effect of CaCl_2_ and NaCl separately on the collagen assembly is known and has been reported before. [47] Analysis of pure collagen hydrogels prepared in the gelling buffers used for the microsphere preparation showed not only a kinetic dependence of the collagen fibrillogenesis (Figure 5A) but also structural differences (Supplementary, Figure S3). Collagen fibers prepared in the presence in GB (10,50) assembled quickly and yielded in a very fine network, giving rise to only a very weak SHIM signal, whilst the presence of GB (50,75) displayed a lag phase prior a fast fibrillogenesis (Figure 5A) and yielded in assembly of a thicker, differently orientated collagen fibers (Figure S3).

SEM images of alginate-collagen microspheres prepared in either of those two chosen gelling buffers and at the absence of additional amino acids such as glycine show significantly different morphologies (Figure S1). Microspheres prepared with 10 mM calcium chloride show a dense surface at the outside and the presence of round structures at the center of the sample (Figure S1A). Distinctive collagen fibers were difficult to identify. It is possible that 10mM CaCl_2_ is not sufficient to fabricate a strong alginate matrix and which, consequently, collapsed into the observed round structures upon preparing the sample for SEM analysis. Previous studies have described a similar phenomenon of a large discrepancy between the alginate morphology of the outer, microsphere, surface and the inner core for samples with an alginate-concentration gradient by both, SEM and size exclusion chromatography. [48] The lack of successfully assembled collagen fibers, as shown in Figure 4B, for this gelling buffer could be a combination of both, the nature of a weak and inhomogeneous alginate network as well as the preference of collagen to assemble to a fine network at the given ratio of calcium, sodium and chloride ions. On the other hand, microspheres prepared with GB (50,75) show a much looser network of alginate for both, outer and inner surface (Figure S1B), indicating the formation of a more stable alginate network due to the increased concentration of calcium ions. Even though occasional round structures can also be found at the center of this sample, indicating the possible collapse of finer structures upon drying, the overall alginate structure is visibly interconnected. Furthermore, collagen fibers can be identified in these images, supporting the observations made in Figure 4B.

## 5 Conclusion

With this work, we were able to show that the *in vitro* collagen fibrillogenesis in alginate microspheres is a complex process which needs a precise analysis in order to yield reproducible IPNs in a spherical geometry. The addition of amino acids increases the amount of collagen fibers assembled in microspheres. This could be because of the shortening of the fibrillogenesis-lag phase. However, the crucial component to consider is the gelling buffer used for preparation and incubation of the microspheres in order to prepare interwoven, porous alginate-collagen IPNs. First clues indicate that the strength of the alginate matrix as well as the ratio of calcium, sodium and chloride ions have a high impact on the formation of collagen fibrils in the gelled or gelling alginate matrix.

## 7 Acknowledgments

The authors thank the Research Council of Norway project number 262893 for financial support under the FRINATEK program. Figure 1 has been created with BioRender.com. The imaging was performed at the Center for Advanced Microscopy, the Norwegian University of Science and Technology (NTNU) with technical assistance from Astrid Bjørkøy. Additionally, the authors would like to thank Mr. Jakob Vinje for the generation of the python-based analysis program for the alginate gelling experiments as well as him and Ms. Jennifer Zehner for their valuable input during the preparation of this manuscript.

