## Supplementary for "Tailoring the assembly of collagen fibers in alginate microspheres"

### Appendix A: Supplementary

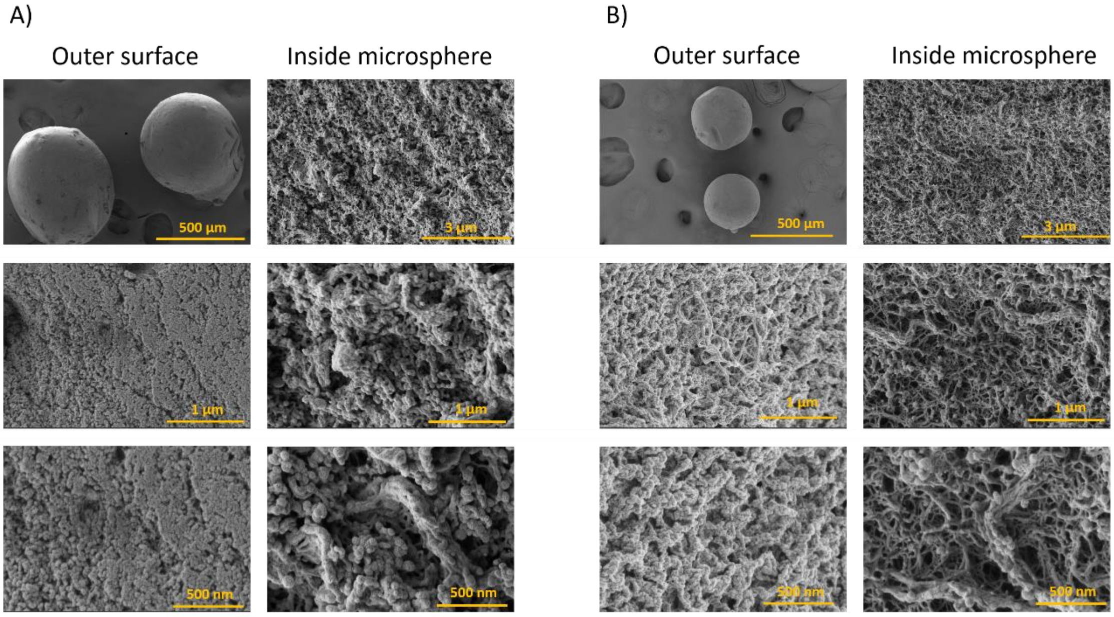


*Figure S1.* *SEM surface analysis of alginate-collagen microspheres. A) Alginate-collagen microspheres prepared in GB (10,50). B) Alginate-collagen microspheres prepared in GB (50,75).*

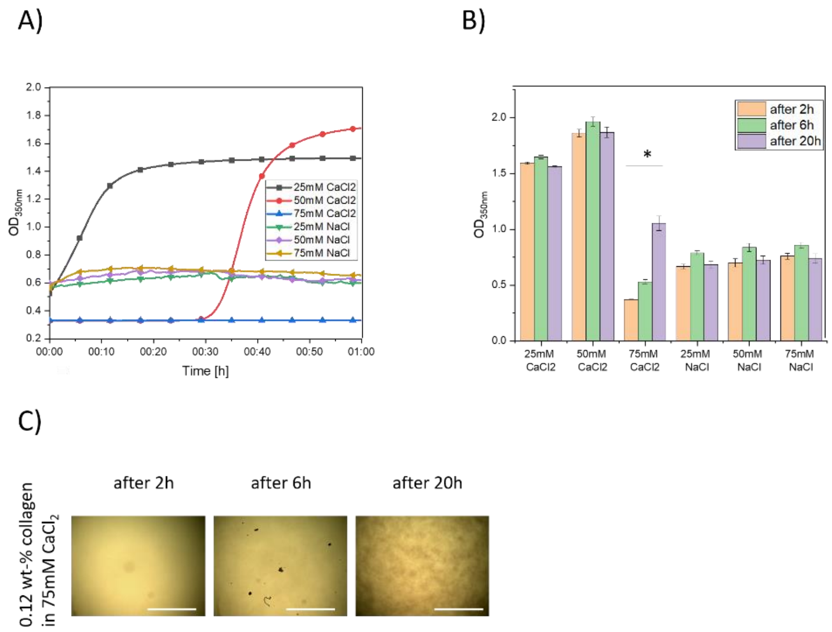


*Figure S2.* *A) Assembly of 0.12 wt-% collagen in solutions containing either NaCl or $CaCl_2$ and 10mM TRIS measured at 37 °C using turbidity assay. B) Measurement of the OD in the respective wells after 2h, 6h and 20h of incubation at 37 °C. In each sample-well the OD has been measured for 9 different locations and the respective value has been averaged. C) Brightfield images of turbidity measurement of collagen assembly for the sample containing 0.12 wt-% collagen and 75mM $CaCl_2$. Images show the slow formation of collagen fibers over time. Scale bar: 1 mm.*

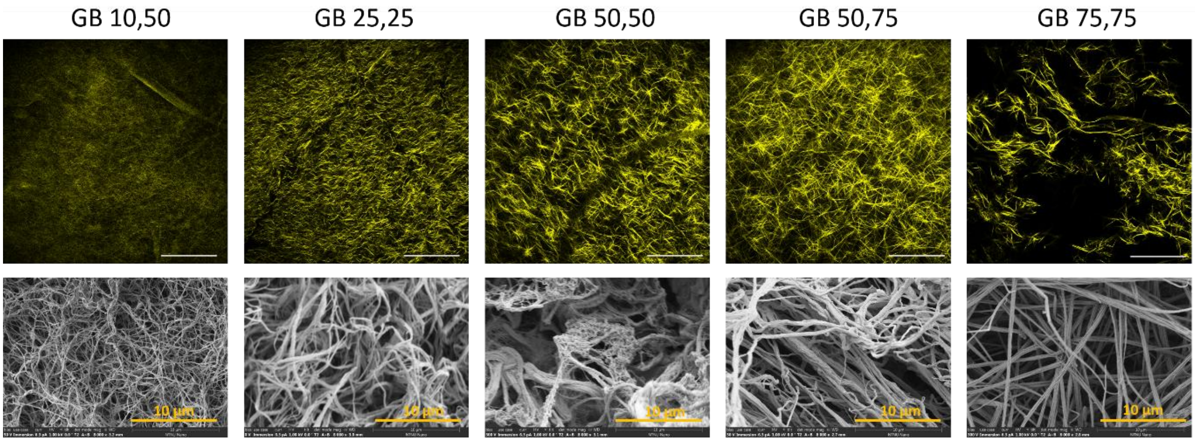


*Figure S3.* *SEM surface analysis and SHIM signals of 0.12 wt-% collagen hydrogels prepared in the presence of the gelling buffers used for the microsphere preparation. Scale bar for SHIM images: 150 µm.*

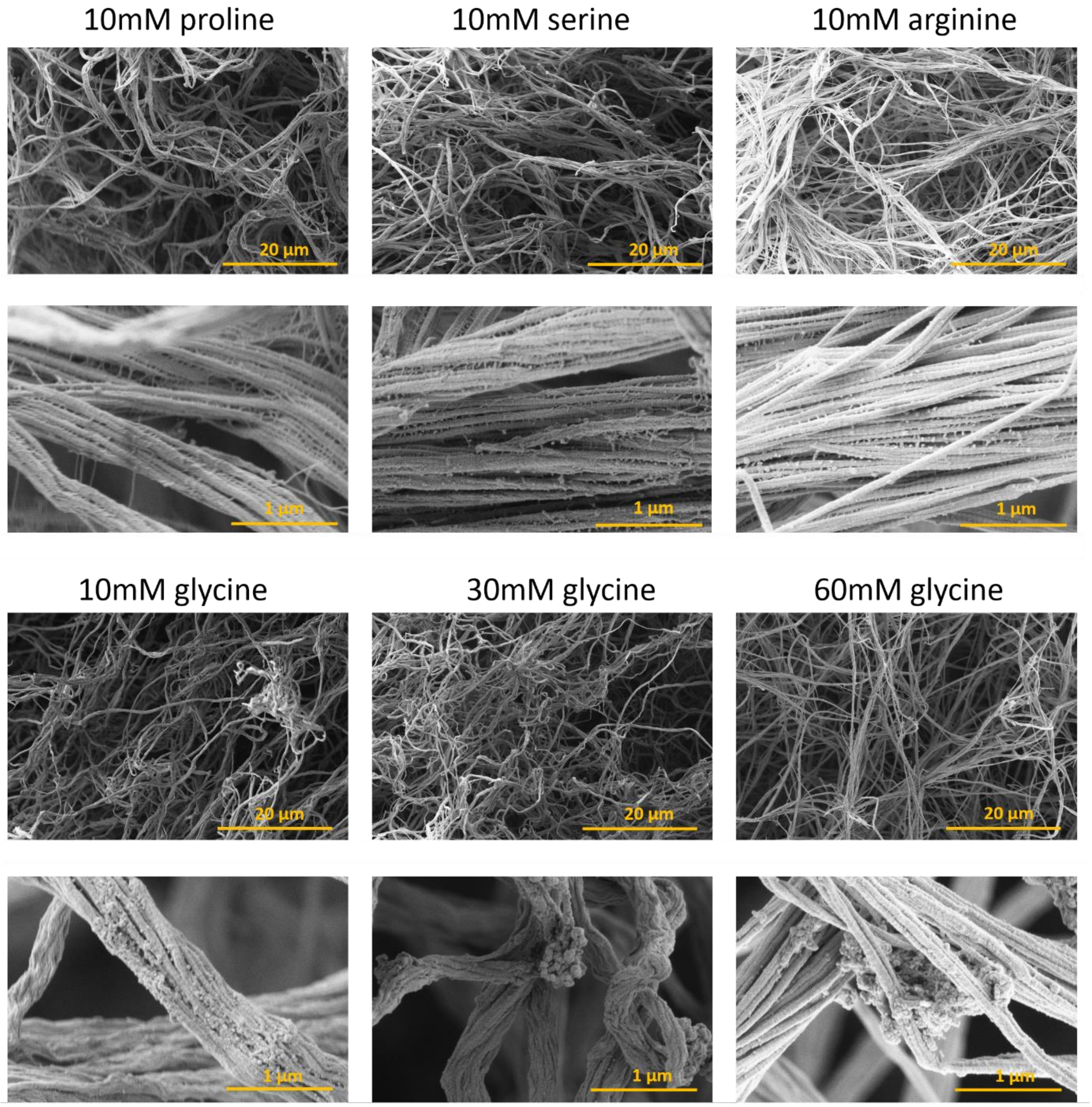


*Figure S4.* *SEM surface analysis of 0.12 wt-% collagen hydrogels prepared with in the presence of amino acids.*

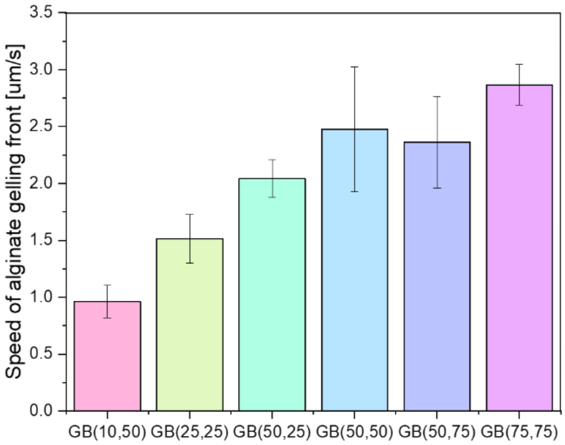


*Figure S5.* *Alginate gelling kinetics of a 1.5 uL 0.6 wt-% alginate droplet in dependence of the used gelling buffer.*
